# plsMD: A plasmid reconstruction tool from short-read assemblies

**DOI:** 10.1101/2025.03.17.643493

**Authors:** Maryam Lotfi, Deena Jalal, Ahmed A Sayed

## Abstract

While whole genome sequencing (WGS) has become a cornerstone of antimicrobial resistance (AMR) surveillance, the reconstruction of plasmid sequences from short-read WGS data remains a challenge due to repetitive sequences and assembly fragmentation. Current computational tools for plasmid identification and binning, such as PlasmidFinder, cBAR, PlasmidSPAdes, and Mob-recon, have limitations in reconstructing full plasmid sequences, hindering downstream analyses like phylogenetic studies and AMR gene tracking. To address this gap, we present plsMD, a tool designed for full plasmid reconstruction from short-read assemblies. plsMD integrates Unicycler assemblies with replicon and full plasmid sequence databases (PlasmidFinder, MOB-typer and PLSDB) to guide plasmid reconstruction through a series of contig manipulations. Using two datasets, one established benchmark dataset used in previous benchmarking studies and another novel dataset consisting of newly sequenced bacterial isolates, plsMD outperformed existing tools in both. In the benchmark dataset, it achieved excellent recall, precision, and F1 scores of 91.3%, 95.5%, and 92.0%, respectively. In the novel dataset, it achieved good recall, precision, and F1 scores of 77.6%, 88.9%, and 74.5%, respectively. plsMD supports two usage modalities: single-sample analysis for plasmid reconstruction and gene annotation, and batch-sample analysis for phylogenetic investigations of plasmid transmission. This computational tool represents a significant advancement in plasmid analysis, offering a robust solution for utilizing existing short-read WGS data to study plasmid-mediated AMR spread and evolution.

**Key points:**

- Accurate plasmid reconstruction from short-read assemblies, surpassing existing binning-based tools.
- Replicon-guided approach enables detection of divergent plasmids.
- Supports single and batch sample analysis, enabling gene annotation as well as plasmid transmission and evolutionary studies.

**Bibliographical note:** The authors are members of the Genomics and Metagenomics Program at Children’s Cancer Hospital Egypt, working in cancer genomics, bioinformatics, and antimicrobial resistance research.

**Graphical abstract:** plsMD uses Unicycler assemblies to identify plasmid replicons via PlasmidFinder and MOB-typer, and aligns the assemblies to PLSDB. It refines alignments, selects reference plasmids, and reconstructs full plasmid sequences. Circular plasmids without replicons are identified separately. plsMD outputs plasmid and non-plasmid FASTA files. Two workflows are supported: single-sample (plasmid/non-plasmid separation, annotation of AMR, VF, IS, and replicons) and batch-sample (grouping plasmids by replicon, MAFFT alignment, rotation, and phylogenetic tree construction). Validation on two datasets showed notably better performance than other benchmarking tools.

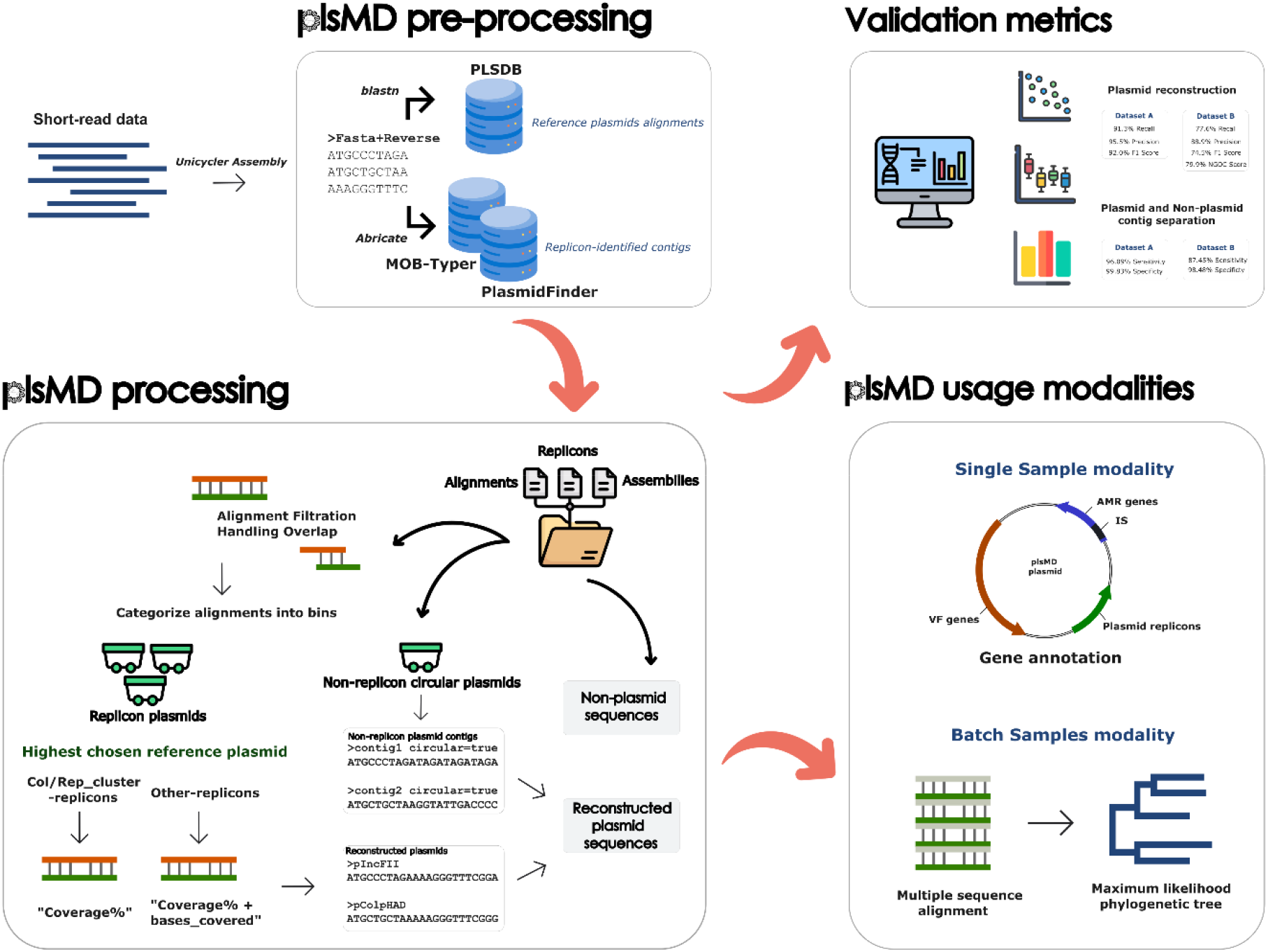

## Introduction

Antimicrobial resistance (AMR) is a critical and growing threat to clinical practice. AMR is primarily disseminated through AMR genes, which are frequently harbored on mobile genetic elements such as plasmids (1). Plasmids are extrachromosomal DNA elements capable of transferring between organisms via mechanisms such as conjugation and transformation. They also replicate independently using specialized sequences called replicons. These replicons have historically served as key markers for plasmid identification and classification (2). Plasmids that share the same replication mechanism are unable to coexist long-term within the same cell (3). As a result, replicon sequences are used to categorize plasmids into distinct incompatibility groups and, labeled with the “Inc” prefix for incompatibility plasmids and the “Col” prefix for colicin-producing plasmids, which encode bactericidal toxins (3,4). Inc-type plasmids tend to be larger in size with several insertion sequences, whereas colicin plasmids are usually smaller and lack repetitive elements (4,5).

Since 2015, the World Health Organization (WHO) has recommended whole genome sequencing (WGS) as a critical tool for AMR surveillance and pathogen detection (6). This has led to its increasing adoption in routine clinical workflows. The vast majority of WGS data is generated using Illumina short-read sequencers, which produce highly accurate reads but face significant challenges during genome assembly (7). In particular, repetitive sequences, commonly found in plasmid DNA, especially near AMR gene cassettes, frequently cause contig breaks during assembly (8). Classical plasmid sequencing approaches rely on the separation of plasmid DNA from chromosomal DNA prior to sequencing. However, these methods struggle to capture large plasmids exceeding 50 kb. To address this, newer methods aim to extract plasmid sequences directly from WGS data (9).

Resolving plasmid sequences from short-read sequencing data has been a persistent challenge (10), yet it remains critically important for understanding AMR gene dissemination and inter-organism gene transfer. Several computational tools have been developed to address this issue. PlasmidFinder relies on a database of replicon sequences to identify and classify contigs as plasmid-derived (9). However, this approach struggles with large plasmids, which often harbor repetitive sequences that cause contig breaks, resulting in only partial plasmid identification (10). cBAR employs a *k*-mer based methodology, using pentamer frequencies to classify contigs as either plasmid or chromosomal, but it does not attempt to bin sequences into distinct plasmid compartments (11). PlasmidSPAdes and Recycler take advantage of assembly graphs derived from De Bruijn graphs (12,13). Recycler searches these graphs for subgraphs that represent distinct plasmids, while PlasmidSPAdes uses depth and circularity characteristics to predict plasmid sequences and group them into bins. Finally, MOB-recon combines in-house databases for complete reference plasmids and plasmid-associated sequences, such as relaxases, to guide the separation of plasmid and chromosomal contigs (14).

While some of these tools perform well in plasmid binning and separating chromosomal contigs, none are designed to reconstruct full contiguous plasmid sequences. This limitation hinders key downstream analyses, such as phylogenetic studies, tracking plasmid evolution, and understanding the mechanisms underlying AMR gene spread. Full plasmid reconstruction with preserved gene order is essential for identifying integron-mediated AMR gene spread, and phylogenetic analysis of complete plasmid constructs is necessary for characterizing plasmid-mediated AMR transmission.

In this study, we present plsMD (https://github.com/Genomics-and-Metagenomics-unit-57357/plsMD), a tool designed for full plasmid reconstruction beyond contig binning from short-read sequence assemblies. This tool utilizes Unicycler assemblies together with established replicon and full plasmid sequence databases to guide plasmid reconstruction. Through a series of contig sequence manipulations, we reconstruct full contiguous plasmid sequences. We evaluated plsMD using two datasets: a benchmark well-characterized dataset from previous studies, and a novel dataset of newly sequenced bacterial isolates. Benchmarking focused on MOB-recon and gplas2, the highest-performing plasmid binning tools currently available. Across both datasets, plsMD outperformed these tools in the accuracy and precision of reconstructed plasmids, as well as in the separation of plasmid and chromosomal contigs. plsMD supports two usage modalities: single-sample analysis, which allows plasmid reconstruction, separation of chromosomal contigs, and subsequent gene annotation using relevant bioinformatics tools; and multi-sample analysis, which groups reconstructed plasmids sharing the same replicons and performs phylogenetic analysis to explore plasmid relatedness and transmission across organisms.

## Materials and Methods

### Annotation and Preprocessing

The workflow for plsMD is summarized in Figure 1. In brief, the input for plsMD consists of assembled shotgun sequencing data using the preferred assembler Unicycler. Other assemblers are compatible, provided they perform circular tagging to match Unicycler’s output. All resulting contigs are reversed and merged with their forward counterparts into a combined FASTA file. plsMD integrates replicon databases; PlasmidFinder (version 18-Apr-2024) and MOB-typer (v3.1.9), and the full plasmid sequences database PLSDB (v. 2024_05_31_v2) respectively (9,15,16).

**Figure 1:**
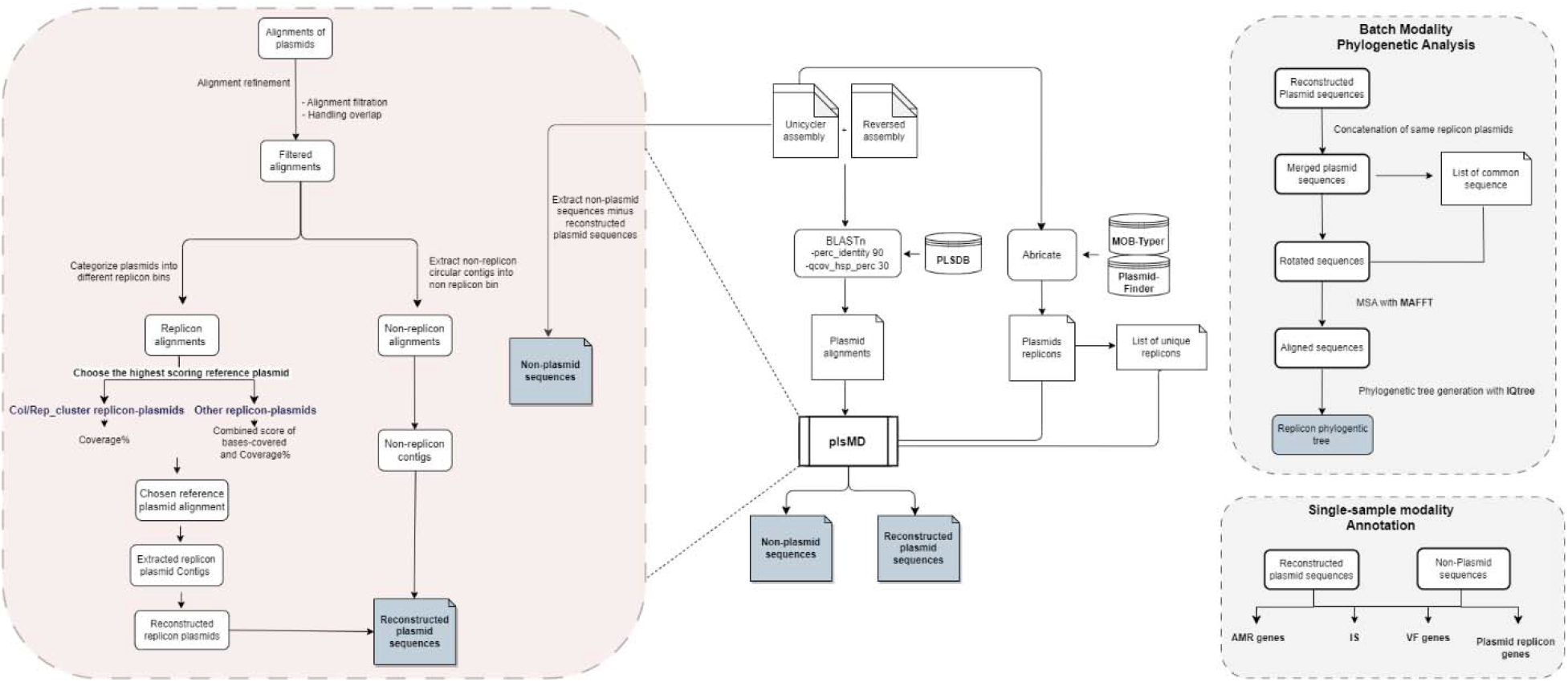
Workflow of the plsMD pipeline. Overview of the plsMD workflow. Unicycler assemblies are used as input and screened for plasmid replicons using PlasmidFinder and MOB-typer. Contigs are aligned against the PLSDB reference database, followed by alignment refinement, overlap handling, and selection of an appropriate reference plasmid based on replicon type. The selected reference plasmid guides reconstruction of a full plasmid sequence. Circular contigs lacking detectable replicons are identified separately. plsMD outputs two FASTA files per sample: reconstructed plasmid sequences and non-plasmid contigs. The tool supports a single-sample workflow for plasmid reconstruction and annotation, and a batch-sample workflow for plasmid grouping, alignment, rotation, and phylogeneti tree construction.

First, Abricate (v 1.0.1, https://github.com/tseemann/abricate) is run on the Unicycler assembly using both the PlasmidFinder and the MOB-typer replicon database to identify plasmid-associated replicons and the contigs they are harbored on. To avoid duplication between the two replicon databases, replicons from MOB-typer are considered for inclusion only if they occur on a different contig than those already identified by PlasmidFinder. The identified replicons serve as anchors for reconstructing plasmid structures from their corresponding contigs. To accommodate cases where same replicon gene exists on multiple contigs (possibly reflecting multiple plasmids sharing the same replicon type), the plasmid names in the Abricate replicon database outputs are modified by appending unique suffixes, allowing the reconstruction of individual plasmid candidates carrying the same replicon. Second, to capture alignments on both forward and reverse orientations, the combined FASTA is aligned to the PLSDB database using BLASTn (17). Parameters are optimized for plasmid detection: - perc_identity 90 and -qcov_hsp_perc 30. The lowered query coverage cutoff accommodates sequence diversity resulting from recombination-based rearrangements, which are common in plasmids. plsMD uses the standard BLAST tabular format (format 6), with two additional fields: query coverage (*qcovs*) and reference plasmid length (*slen*). For clarity, “contig” is used to refer to the BLAST query sequence and “reference plasmid” to the BLAST subject sequence throughout this section.

### Alignment Refinement

#### Filtration

To ensure all plasmids are reconstructed in the correct (reference) plasmid orientation, alignments mapping in reverse on the reference plasmid are removed, as the reverse-orientation contigs generated during preprocessing inherently cover this direction. Contigs annotated as circular in the original unicycler assembly are flagged. If a reference plasmid is aligned by both circular and non-circular contigs, it is excluded entirely to preclude the merging of biologically distinct forms.

#### Handling overlap

plsMD detects and resolves nested, partially overlapping and repeated contig alignments to prevent redundant sequence incorporation. (i) Nested contigs alignments: Alignments fully contained within the alignment interval of a larger contig are removed as they provide no additional sequence information. These nested contigs are also excluded from the final non-plasmid set. (ii) Partially overlapping contig alignments; are processed by calculating the overlap between contigs using reference plasmid alignment coordinates and the number of overlapped bases to be used for trimming from contigs during reconstruction. (iii) Repeated contig alignments within the same reference plasmid; the alignments are retained only if the alignment involves more that 80% of the contig. This high cutoff ensures inclusion of only substantially covered repetitions.

### Reference Plasmid identification for reconstruction

#### Bases covered and coverage Calculations

*Bases covered* is calculated as the merged non-overlapping alignment intervals per reference plasmids. *Coverage percentage* is calculated as the proportion of the length of the reference plasmid spanned by bases covered:

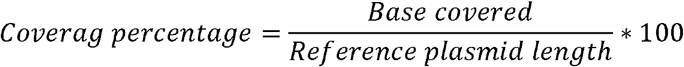

#### Reference plasmid choice

Bins are created for each replicon in a sample, with only plasmid alignments containing a specific replicon contig binned together for reference plasmid selection. To choose the highest scoring reference plasmid for each reconstructed plasmid, plsMD uses two different scoring approaches depending on the replicon type: ***(i) Col and rep_cluster replicons;*** The reference plasmid is chosen based on the highest coverage percentage as these replicon plasmids are typically smaller and have few or no insertion sequences; ***(ii) Inc/Other/Col-hybrid replicons;*** Reference plasmid choice is based on a average scoring of both coverage percentage and bases covered, to allow larger plasmid capturing as these plasmids commonly contain repetitive elements. For Inc/Other/Col-hybrid replicons plasmids, to prioritize reference plasmids with high coverage, a progressive coverage filter is applied, starting at 80% and decreasing in 10% increments if no plasmids meet the threshold (with no lower bound), ensuring the detection of all possible replicon plasmids.

#### Redundant plasmid removal

All chosen reference plasmids across all replicons are checked for redundancy as follows; (i) If the same reference plasmid is chosen across different replicon bins, such as if a plasmid has multiple replicons, only one instance is included in the final reconstructed plasmid file for each sample. (ii) For hybrid Col-Inc plasmids, the chosen plasmid in the Col replicon bin is excluded, as the selected Inc plasmid would be more comprehensive in its selection of reference plasmid.

### plsMD output sequences

#### Plasmid sequences

##### (i) Reconstructed Replicon plasmids

Reconstruction of the plasmid is done by extracting the contigs for the chosen plasmid from the combined FASTA files. Overlapping regions in the contigs are trimmed, and the contigs are merged into a single sequence for each plasmid. Naming of the reconstructed plasmid is based on the created replicon bins.

##### (ii) Non-replicon plasmids

Contigs marked for circularity, that do not harbor any replicons are added as such in the reconstructed plasmid file and named as a circular contig.

#### Non-plasmid sequences

Contigs included in plasmid sequences are excluded from assembled FASTA files to generate non-plasmid contigs. Additionally, nested contigs that align to reference plasmids but were excluded from plasmid assembly due to redundancy are also removed from the non-plasmid contigs.

### Usage Modalities

#### Single Sample modality – Annotation

The separation of plasmid and non-plasmid sequences enables the annotation of various genetic elements, including AMR genes, virulence factor (VF) genes, insertion sequences, and plasmid replicons. Annotation is performed using Abricate and AMRFinder (18) for AMR gene identification, VFDB (19) for virulence factor genes, ISfinder (20) for insertion sequences, and PlasmidFinder (9) for plasmid replicons.

#### Batch modality – Phylogenetic Analysis

For the purpose of investigating phylogenetic relationships between identified plasmids in different samples, reconstructed plasmids from different samples with same replicons (in the same replicon bin) are grouped together in one multi-fasta file. Prior to multiple sequence alignment, plasmid sequences must be rotated to a common start point to resolve inconsistencies caused by arbitrary starting positions in circular plasmid sequences. This is achieved by identifying a 10-30 bp shared sequence between the plasmids, and adjusting plasmid sequences to start from this point, reversing them if necessary. Once rotated, the sequences are aligned using MAFFT v7.525 (21), which automatically determines the optimal alignment strategy. Finally, a maximum likelihood phylogenetic tree is generated using IQ-TREE v2.3.6 and visualized in iTOL version 7 (22).

### Validation

#### Dataset Validation and Quality Control

To evaluate plsMD, we compared it with two commonly used plasmid binning tools, Mob-recon (14) and gplas2 (23), using the benchmark dataset compiled by Robertson and Nash (14). This benchmark dataset originally consisted of 133 closed genomes and 377 associated plasmids sequenced using both PacBio and Illumina technologies. The Illumina short-reads for these samples were assembled using Unicycler (v0.5.0) with default parameters. To ensure data integrity and completeness, we implemented stringent validation criteria by blasting assembled contigs to complete reference genomes from the same samples with 90% identity cutoff. Entire samples were excluded if any complete genome showed less than 80% collective coverage by all contigs, indicating potential mislabeling or contamination. Additionally, samples were excluded if any contig longer than 1000 bp aligned to a complete genome with less than 80% of its length, ensuring all substantial contigs were fully explained by the provided complete genomes and preventing inclusion of samples with incomplete reference data. Applying these rigorous criteria resulted in 80 high-quality samples with 244 plasmids being included in our validation, herein called the filtered benchmark dataset.

#### Independent Validation with Novel Plasmids

To address potential database dependency and demonstrate plsMD’s performance with novel plasmids, we additionally validated the tool on a dataset comprising samples sequenced from June 2024 onward, confirmed absent from PLSDB (v. 2024_05_31_v2). This independent validation included 68 samples with 269 plasmids that were not present in our reference databases, herein called the novel dataset. Details of both datasets regarding the species composition, plasmid size distribution, and replicon types for both datasets are presented in Supplementary Table 1 and summarized in Table 1.

**Table 1:**
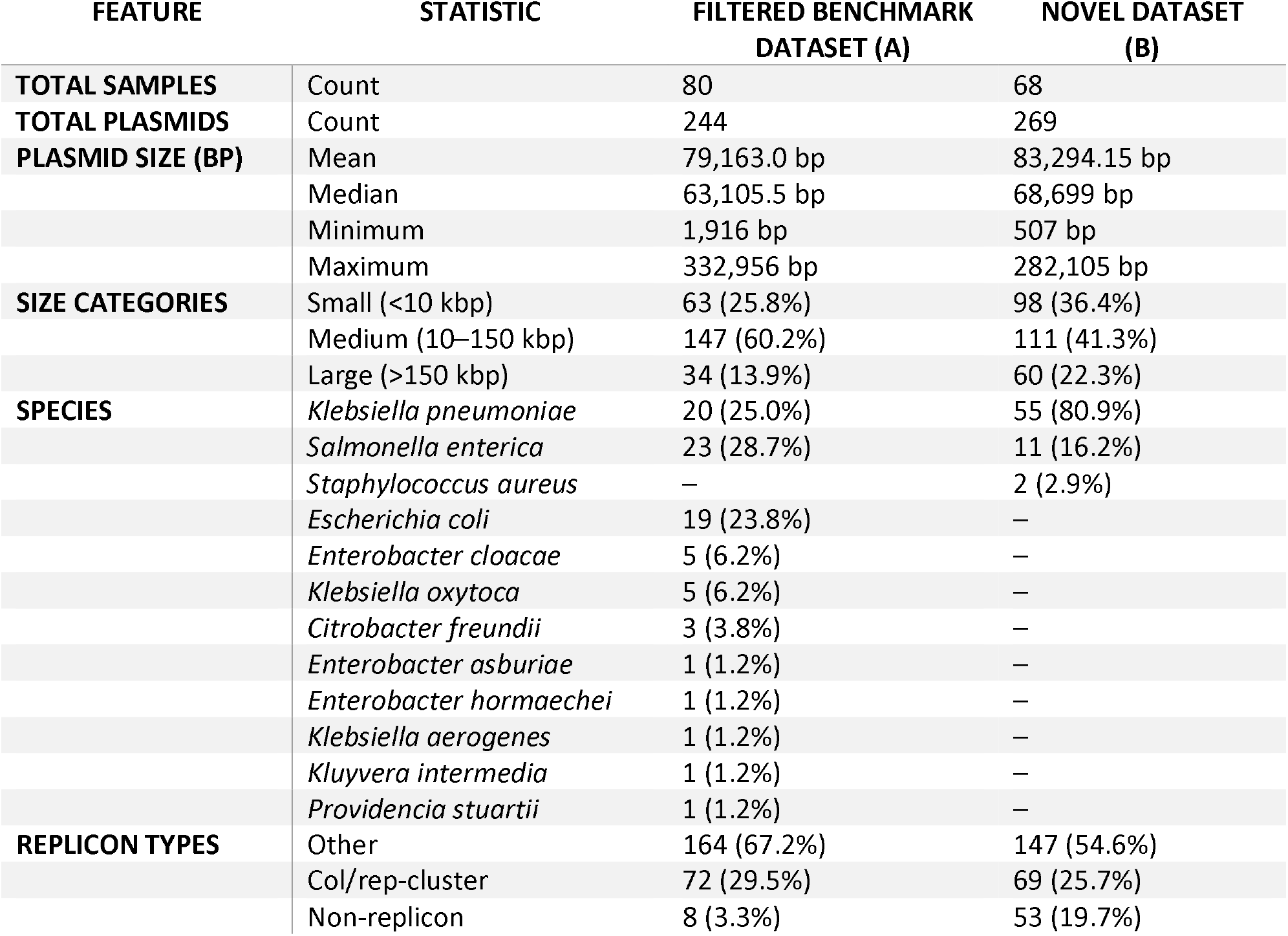
Description of the datasets used in the benchmarking studies.

### Evaluation metrics

#### Plasmid Reconstruction Assessment: Recall and Precision

For each sample, each reconstructed plasmid was separately aligned against all the ground truth plasmids. The recall metric evaluates the alignment relative to the ground truth plasmid. First *the ground truth plasmid length* is calculated. Then, *bases covered* is calculated as the merged non-overlapping alignment intervals per ground truth plasmid. And finally, *coverage percentage* is calculated as the proportion of the length of the ground truth plasmid spanned by *bases covered*. The precision metric, on the other hand, evaluates the alignment relative to the reconstructed plasmid. The *reconstructed plasmid length* is calculated as the total length of the reconstructed plasmid sequence. For MOB-recon and gplas2, which generate multi-FASTA plasmid bins without contig duplication, reconstructed plasmid length accounts for repeated contigs if they align multiple times to the same ground truth plasmid. Then, *aligned length* is calculated as the merged non-overlapping alignment intervals per reconstructed plasmid. Finally, *query coverage* is calculated as the proportion of the length of the reconstructed plasmid spanned by *aligned length*.

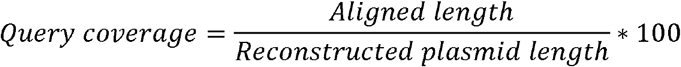

Pairs of reconstructed plasmid and ground truth plasmid alignments were extracted along with their associated metrics: *bases covered, coverage percentage, aligned length*, and *query coverage*. A Hungarian algorithm was implemented to find the optimal matching between reconstructed plasmid and ground truth plasmid sequences, ensuring that each reconstructed plasmid and ground truth plasmid were matched only once. The algorithm maximizes the total *coverage percentage* (recall) and query coverage (precision) by assigning each query to its best-matching subject. This is achieved by constructing a weight matrix based on the sum of both values. The recall of each plasmid was represented by the *coverage percentage* chosen by the matching algorithm, while the precision of each reconstructed plasmid was determined by the *query coverage* assigned by the algorithm. Reconstructed plasmid sequences without a match were assigned *query coverage*= 0, while ground truth plasmid sequences without a match were assigned *coverage percentage* = 0.

Average recall for each tool was calculated as the mean *coverage percentage* across all ground truth plasmids, while average precision was computed as the mean *query coverage* across all reconstructed plasmids for each tool.

### Plasmid Contig Classification: Sensitivity and Specificity

For each sample, all reconstructed plasmids and non-plasmid contigs were aligned against the corresponding ground truth plasmid and chromosome sequences. The *aligned length* from aligning reconstructed plasmids against ground truth plasmid sequences represented True Positives (TP), while alignment against ground truth chromosome sequences represented False Positives (FP). Similarly, *aligned length* from aligning non-plasmid sequences against ground truth plasmid sequences represented False Negatives (FN), while alignment against ground truth chromosome sequences represented True Negatives (TN). Sensitivity and Specificity were calculated as follows:

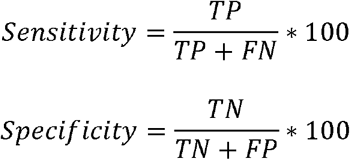

#### Normalized Gene Order Conservation (NGOC) score calculation

To assess syntenic fidelity between reconstructed and ground truth plasmids, a Normalized Gene Order Conservation (NGOC) score was calculated. Reconstructed plasmids and corresponding ground truth plasmids were annotated using Prokka (v1.14.5) (24). Gene order information was extracted from the resulting GFF files, and all adjacent gene pairs were generated for each plasmid, accounting for plasmid circularity and bidirectional gene orientation. The NGOC score was defined as the proportion of adjacent gene pairs in the reference plasmid that were conserved in the reconstructed plasmid. The relationship between NGOC score and plasmid recall percentage was evaluated using Pearson correlation.

#### Statistical analysis

Differences in recall between tools were assessed using the Wilcoxon signed-rank test for paired comparisons. To account for multiple comparisons, p-values were adjusted using the Benjamini-Hochberg false discovery rate (FDR) procedure. An FDR-adjusted p ≤ 0.05 was considered statistically significant.

## Results

### Plasmid Reconstruction Performance

We compared the performance of plsMD with the two established plasmid binning tools; MOB-recon and gplas2, assessing reconstruction accuracy using recall (completeness) and precision (accuracy) (Supplementary Table 2 and Figure 2a and d). On the filtered benchmark dataset (n=244 plasmids), plsMD achieved the highest overall recall (93.07%), substantially greater than MOB-recon (85.97%) and gplas2 (75.2%). This high level of completeness was supported by the number of fully failed reconstructions (recall =0): plsMD failed to reconstruct 7 plasmids, whereas MOB-recon and gplas2 failed to reconstruct 9 plasmids and 30 plasmids, respectively. Furthermore, plsMD demonstrated superior precision (95.47%) compared to MOB-recon (93.02%) and gplas2 (85.77%). This high precision was evidenced by the number of spurious assemblies (precision =0): plsMD produced only 3 extra plasmids, compared to 9 for MOB-recon and 32 for gplas2. When performance was stratified by plasmid size and replicon type (Figure 2b and c), plsMD maintained significantly higher recall than MOB-recon and gplas2 across all plasmid size categories—small (<10 kbp), medium (10–50 kbp), and large (>50 kbp) (Wilcoxin test p < 0.05). Stratified by replicon type, plsMD scored the lowest recall amongst non-replicon plasmids, but exhibited higher recall than both MOB-recon and gplas2 in both Col/rep-cluster and Other replicon types. This combined superior performance is reflected in its high F1 score of 91.94%, which significantly outperformed MOB-recon (84.93%) and gplas2 (68.86%).

**Figure 2:**
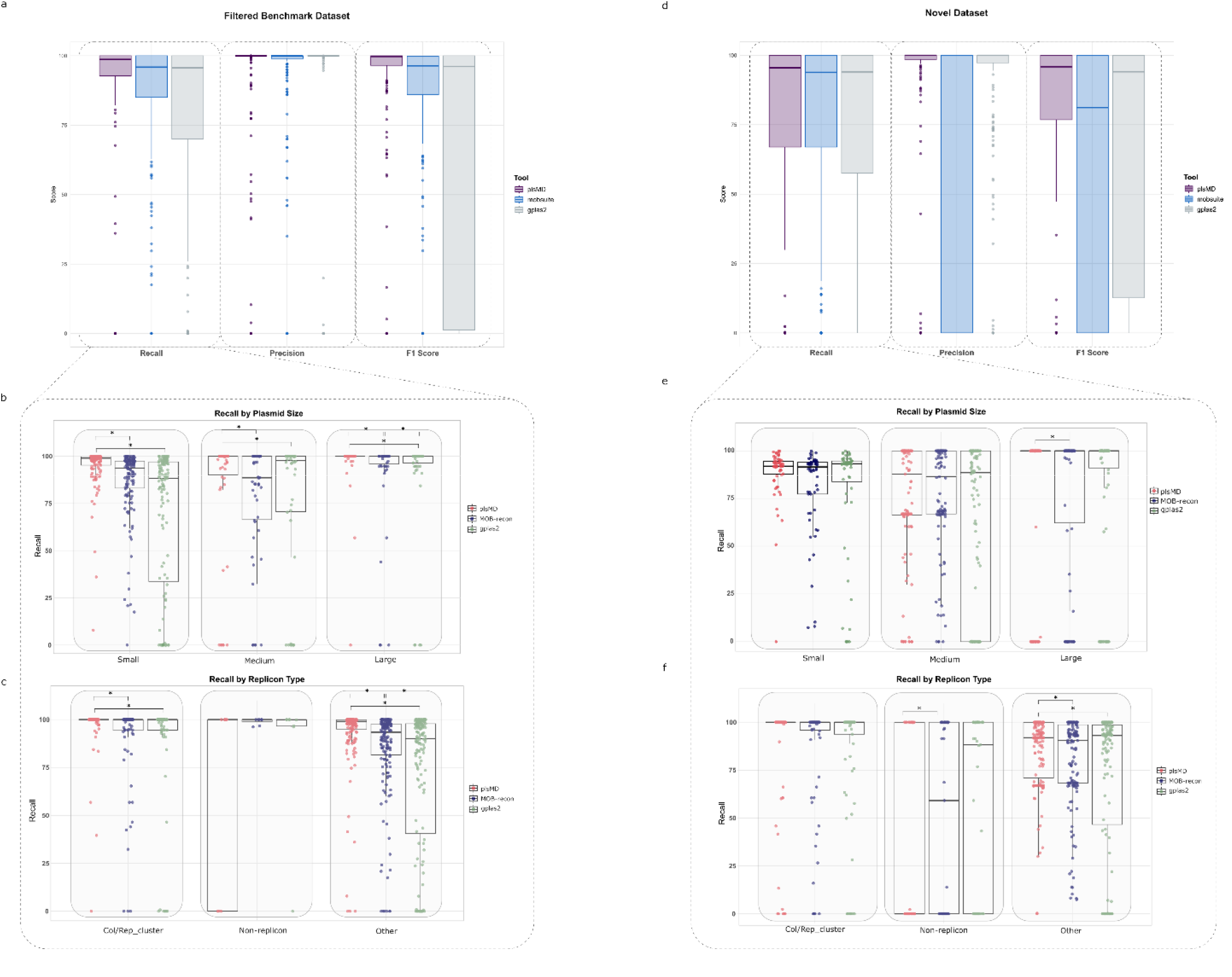
Benchmarking performance of plsMD compared with plasmid binning tools. Comparison of plsMD with MOB-recon and gplas2. (a) Recall, precision, and F1 score for all plasmids in the filtered benchmark dataset (n = 244). (b) Recall stratified by plasmid size category (small <10 kbp, medium 10– 50 kbp, large >50 kbp) in the filtered benchmark dataset. (c) Recall stratified by plasmid replicon category (Col/rep-cluster, other, and non-replicon plasmids) in the filtered benchmark dataset. (d) Recall, precision, and F1 score for all plasmids in the novel dataset (n = 269). (e) Recall stratified by plasmid size category in the novel dataset. (f) Recall stratified by plasmid replicon category in the novel dataset. Statistical significance was assessed using paired Wilcoxon signed-rank tests with false discovery rate (FDR) correction.

To assess robustness against database dependency, we conducted an independent validation on a Novel Dataset (n=269 plasmids) confirmed to be absent from our PLSDB reference (Figure 2d). plsMD achieved the highest overall recall (77.61%) compared to MOB-recon (76.84%) and gplas2 (72.77%). In this dataset, plsMD failed to reconstruct 34 plasmids, compared to 25 for MOB-recon and 53 for gplas2. For precision, plsMD again outperformed the other tools (88.89%) compared to MOB-recon (70.01%) and gplas2 (87.57%). This accuracy was supported as plsMD produced only 19 extra plasmids compared to 89 for MOB-recon and 14 for gplas2. Crucially, recall by plsMD remained significantly higher than MOB-recon for large (>50 kbp) plasmids (Wilcoxin test p < 0.05), a strength reflected in its significantly higher recall for plasmids with Other replicon types (which are frequently larger), although it scored lower recall than gplas2 amongst non-replicon plasmids (Figure 2e and f). Even with novel sequences, plsMD maintained the highest overall F1 score of 74.5%, surpassing MOB-recon (59.20%) and gplas2 (68.65%).

### Plasmid and Chromosomal Contig Separation

For assessing contig classification to plasmid and non-plasmid compartments, we calculated sensitivity (percentage of correctly classified plasmid contigs) and specificity (percentage of correctly classified non-plasmid contigs) for both datasets (Supplementary Table 3). All three tools demonstrated proficiency in basic sequence partitioning, showing high and comparable contig classification accuracy across both datasets. In the Filtered Benchmark Dataset, plsMD scored 96.09% sensitivity and 99.83% specificity, closely matching MOB-recon’s 96.05% sensitivity and 99.84% specificity, while gplas2 achieved 93.02% sensitivity and 98.58% specificity. In the Novel Dataset, plsMD achieved 87.45% sensitivity and 98.48% specificity. This performance was slightly lower than the other tools on the novel set, where MOB-recon scored 93.02% sensitivity and 99.81% specificity, and gplas2 scored 93.08% sensitivity and 99.9% specificity.

### Syntenic preservation by plsMD

The core innovation of plsMD lies in its ability to generate contiguous, full plasmid sequences – an outcome not achieved by plasmid binning tools. To validate the quality and order of the genes within these assemblies, we calculated the Normalized Gene Order Conservation (NGOC) score, a metric for syntenic fidelity. We report an average NGOC score of 79.9% (Figure 3a), demonstrating the high fidelity of plsMD assemblies in preserving gene context. This accuracy is further supported by a strong correlation between recall and the NGOC score (Figure 3b), confirming that when a plasmid is properly assembled (high recall), it is also assembled with the correct gene order. An example comparison between a ground truth plasmid and its plsMD reconstruction confirms conserved gene order (Figure 3c).

**Figure 3:**
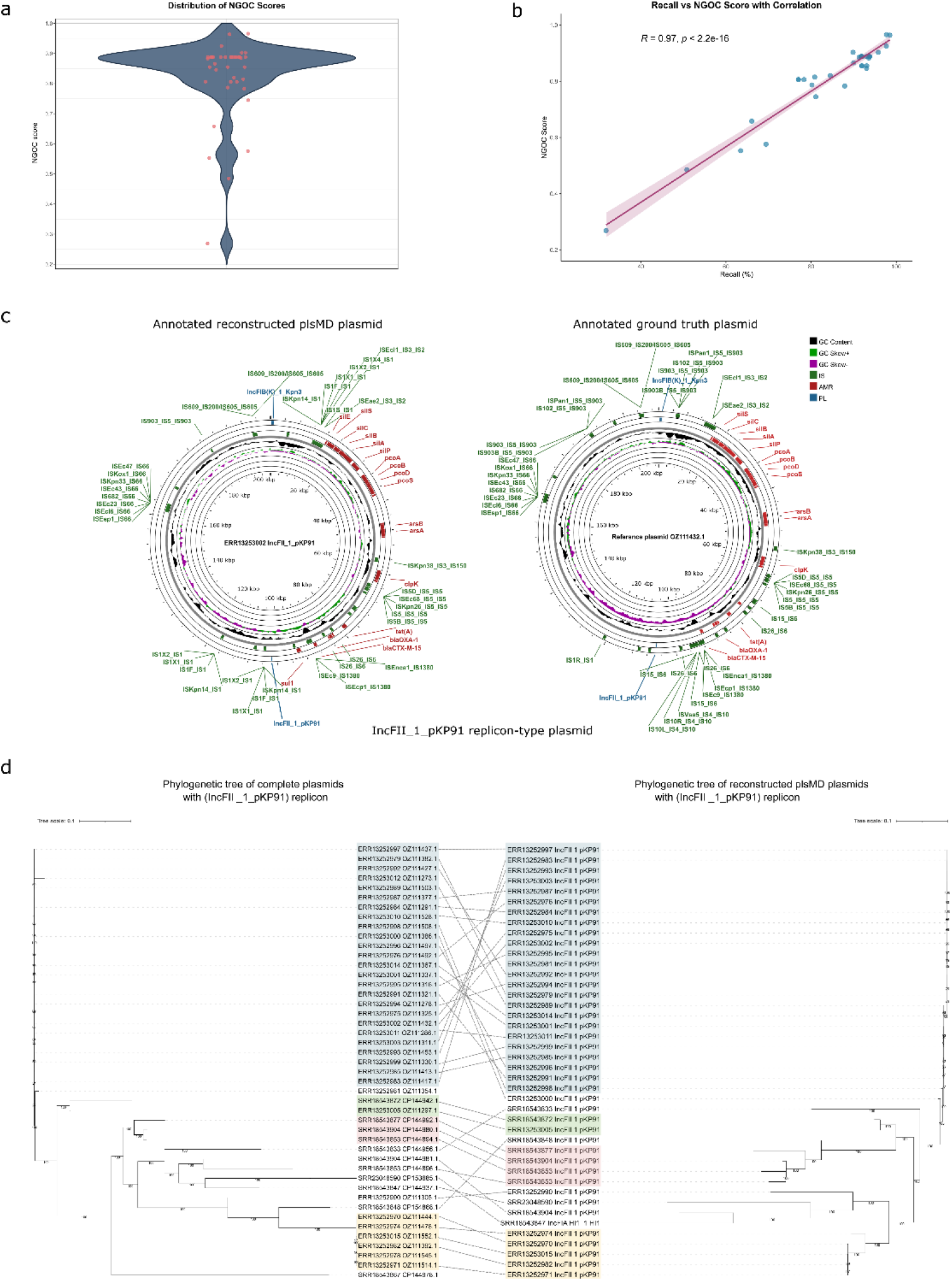
Gene order conservation and phylogenetic consistency of plsMD reconstructions. (a) Normalized Gene Order Conservation (NGOC) scores for plasmids reconstructed using plsMD relative to their ground truth counterparts, reflecting preservation of gene order. (b) Correlation between NGOC score and plasmid recall, calculated using Pearson correlation. (c) Example comparison between a full closed ground truth plasmid harboring the IncFII(pKP91) replicon and its corresponding plsMD reconstruction from the same sample. Sequences are annotated for plasmid replicons, antimicrobial resistance (AMR) genes, virulence factor (VF) genes, and insertion sequences (ISs). (d) Maximum-likelihood phylogenetic trees constructed from complete ground truth plasmids harboring the IncFII(pKP91) replicon and from the corresponding plsMD-reconstructed plasmids. Trees were inferred using IQ-TREE and visualized using iTOL; conserved clustering patterns are highlighted.

### Applicability of plsMD for plasmid phylogenetic studies

Tracking and transmission analysis of AMR gene spread is increasingly required. Since the majority of AMR genes are harbored on plasmids, accurate plasmid phylogenetic analysis is essential, and this process is supported by the batch mode of plsMD. To test the accuracy of plsMD reconstructions for this downstream application, we compared phylogenetic trees built from reconstructed plasmids to those built from ground truth plasmids. We used plasmids harboring the IncFII(pKP91) replicon from the Novel Dataset as our test case, as it had high prevalence across 45 samples. The phylogenetic trees generated for both the complete ground truth plasmids harboring the IncFII(pKP91) replicon as well as those reconstructed from plsMD are visualized using iTOL version 7 (Figure 3d). The conserved clustering of plasmids from both trees demonstrates the high fidelity of plsMD reconstructions for use in transmission and evolutionary studies.

## Discussion

With the rapid advancements in long-read sequencing technologies and the development of newer sequencing techniques, the future of plasmid identification and sequencing is promising. However, for the vast amount of existing short-read sequencing data, current tools have yet to achieve the precision and reliability needed to accurately extract plasmid sequences. Plasmid identification and binning tools generally rely on three main approaches. Assembly graph inspection methods, such as PlasmidSPAdes and Recycler, search for sub-graphs within assembly graphs. However, these tools typically score low on precision and recall and often fail to classify large plasmids with highly repetitive sequences due to the fragmented and ambiguous assembly graphs they produce (10). Reference-based approaches, including PlasmidFinder and MOB-recon, depend on precompiled databases for plasmid identification. While these methods can be effective, they inherently struggle to detect novel plasmids, limiting their broader applicability. Algorithm-based methods, such as cBAR (11) and PlasmidHunter (25), use sequence-specific features (e.g., GC content or *k*-mer frequencies) to identify plasmids. These approaches perform well in metagenomic contexts but generally do not attempt plasmid binning, as they cannot fully resolve sequences from different plasmids based solely on sequence characteristics (11,25). Hybrid approaches, such as PlasBin-flow and gplas2, attempt to combine these methods to improve performance (26).

plsMD, while being a reference-based tool, operates on the premise that for a plasmid to be effectively maintained across cell generations, it requires independent replication, guiding the reconstruction of plasmids around replicon genes. This approach facilitates the identification of highly divergent plasmids by allowing reconstruction even with low similarity to reference plasmids. Furthermore, this replicon-guided approach provides a stronger anchor for plasmid construction compared to methods like MOB-recon, which use other plasmid-associated genes such as relaxases. This difference is believed to contribute to plsMD’s higher precision and lower number of extra plasmids. Additionally, the use of PLSDB as a reference database, which comprises complete plasmid sequences from diverse bacterial species (27), avoids organism-specific constraints during reconstruction.

Consequently, while the performance of plsMD is inherently influenced by the quality of sequencing reads, assembly accuracy, and the comprehensiveness of the reference database, it achieves superior recall and precision compared to existing tools, with fewer instances of non-reconstructed and extra plasmids. To accomplish this across varying plasmid sizes, plsMD employs a non-uniform strategy. Specifically, recognizing that repetitive sequences often cause contig breaks in Illumina-based assemblies, plsMD distinguishes between low-repetitive sequence plasmids, such as Col replicon plasmids, and those with highly repetitive sequences. This enables the complete and accurate assembly of plasmids across a wide size range.

Despite the highly accurate reconstruction achieved by plsMD, its methodology presents certain inherent limitations. The dependence of plsMD on the initial detection of replicon genes, causes the tool to struggle with non-replicon plasmids (those without detectable replicons in the current databases), resulting in lower recall for this category in benchmarking. The ability of plsMD to capture non-replicon sequences is therefore limited, primarily relying on the contig being tagged as circular. An additional, minor limitation is that the process of preventing redundant sequence incorporation during assembly may cause small mobile elements or integrated plasmid segments to be excluded from the final non-plasmid (chromosomal) output, slightly impacting the completeness of that file. We view this as a minor trade-off, however, as achieving 100% complete and contiguous chromosomal sequences from short-read assembly alone often necessitates the use of specialized hybrid assembly or reference-based finishing tools, minimizing the practical impact of this specific omission.

The initial development of plsMD was motivated by the need to generate full contiguous plasmid sequences from short-read data for evolutionary and transmission studies. In this context, the information provided by full contiguous sequences is pivotal for plasmid phylogenetic analysis, which uncovers hidden transmission routes often overlooked when focusing solely on clonal expansion (28). The highly accurate reconstruction of plsMD with conserved gene order provides strong advantages for plasmid evolutionary studies. Furthermore, accurate gene order conservation provides further information regarding integron-mediated recombination events, which play major roles in AMR co-resistance and evolution.

Overall, plsMD uses established databases and tools in a streamlined workflow to address a critical and challenging problem without relying on complex algorithms or methodologies. While plasmid binning may be sufficient in a range of applications, plsMD provides fully assembled and polished plasmid sequences, opening up opportunities for a broad range of analyses and insights that otherwise would not be possible.

## Conclusion

In this study, we present plsMD, a computational tool designed to reconstruct full, contiguous plasmid sequences from short-read whole genome assemblies. By combining replicon-guided reconstruction with reference plasmid alignment, plsMD overcomes key limitations of existing plasmid binning approaches, particularly for large and repetitive plasmids. Benchmarking against state-of-the-art tools demonstrated that plsMD achieves higher recall, precision, and improved preservation of gene order across both well-characterized and novel plasmid datasets.

Importantly, plsMD enables downstream analyses that require complete plasmid sequences, including plasmid phylogenetics, transmission tracking, and investigation of integron-mediated antimicrobial resistance evolution. By allowing accurate plasmid reconstruction from widely available Illumina sequencing data, plsMD provides a practical and robust solution for plasmid-focused genomic studies and supports improved surveillance of plasmid-mediated antimicrobial resistance.

## Declarations

### Consent to Participate

Not applicable.

### Consent to Publish

Not applicable.

### Ethics declaration

Not applicable.

### Author Contribution

D.J. conceptualized the study. D.J. and M.L. developed the methodology and wrote the manuscript. M.L. prepared all figures. A.S. provided supervision and reviewed the manuscript. All authors read and approved the final manuscript.

### Availability and Requirements

Project name: plsMD

Project home page: https://github.com/Genomics-and-Metagenomics-Unit-57357/plsMD

Operating system(s): Linux, macOS

Programming language: Bash, Python 3

Other requirements: plsMD integrates and internally executes the following third-party tools as part of its automated workflow: Unicycler, BLAST+, Abricate, MAFFT, and IQ-TREE. These dependencies are handled within the plsMD pipeline and do not require independent execution by the user. Python 3 with standard scientific Python libraries is required to run plsMD.

License: MIT License

Any restrictions to use by non-academics: None

### Availability of Data and Materials

The benchmark dataset used in this study was obtained from the publicly available dataset published by Robertson and Nash as part of the MOB-suite study (14). This dataset comprises Illumina short-read assemblies and corresponding closed reference genomes and plasmids generated using long-read sequencing. The novel dataset consists of bacterial isolates sequenced as part of this study from June 2024 onward and was confirmed to be absent from the PLSDB reference database (version 2024_05_31_v2) at the time of analysis. Details of samples included in both datasets are presented in Supplementary Table 1. The plsMD source code and documentation are available at https://github.com/Genomics-and-Metagenomics-Unit-57357/plsMD.

